# Isolation of lactic acid bacteria capable of reducing environmental alkyl and fatty acid hydroperoxides, and the effect of their oral administration on oxidative-stressed nematodes and rats

**DOI:** 10.1101/592162

**Authors:** Akio Watanabe, Takuro Yamaguchi, Kaeko Murota, Tadaaki Ishii, Junji Terao, Sanae Okada, Naoto Tanaka, Shinya Kimata, Akira Abe, Tomonori Suzuki, Masataka Uchino, Youichi Niimura

**Author notes:** Institute of Agricultural and Life Sciences, Academic Assembly, Shimane University, Matsue, Simane 690-8504 Japan (Kaeko Murota). Faculty of Clinical Nutrition and Dietetics, Konan Women’s University, Higashinada-ku, Kobe, Hyogo 658-0001 Japan (Junji Terao).

## Abstract

Reinforcement of hydroperoxide-eliminating activity in the intestines and colon should prevent associated diseases. We previously isolated a lactic acid bacterium, *Pediococcus pentosaceus* Be1, that facilitates a 2-electron reduction of hydrogen peroxide to water. In this study, we successfully isolated an alternative lactic acid bacterium, *Lactobacillus plantarum* P1-2, that can efficiently reduce environmental alkyl hydroperoxides and fatty acid hydroperoxides to their corresponding hydroxy derivatives through a 2-electron reduction. Each strain exhibited a wide concentration range with regard to the environmental reducing activity for each hydroperoxide. Given this, the two lactic acid bacteria were orally administered to the oxygen-sensitive short-lived nematode mutant, and this resulted in a significant expansion of its lifespan. This observation suggests that *P. pentosaceus* Be1 and *L. plantarum* P1-2 inhibit internal oxidative stress. To determine the specific organs involved in this response, we performed a similar experiment in rats, involving induced lipid peroxidation by iron-overloading. We observed that only *L. plantarum* P1-2 inhibited colonic mucosa lipid peroxidation in rats with induced oxidative stress.

## Introduction

The intestines and colon are key points where defense mechanisms against various types of diseases and stresses are employed. Such diseases are often triggered by oxidative stress, which is the primary stress at these points *in vivo*. Although hydroperoxides (i.e., hydrogen peroxide and lipid hydroperoxide) are major causes of oxidative stress, the reductase activity in colonic mucosa for hydroperoxides is lower than that in other organ tissues [1]. Enhancement of hydroperoxide reductase activity in the colonic mucosa can prevent bowel diseases. It has been shown that administration of *Lactococcus lactis*, which produces a catalase from the *Bacillus* gene, prevents chemically induced colon cancer in mice [2]. Similar to hydrogen peroxide, lipid hydroperoxide is a downstream reaction product of ROS that strongly contributes to bowel disease [3]. A number of chemical anti-oxidant treatments for lipid hydroperoxide exist [4-6]. The microbial antioxidative effects of the *Streptococcus thermophilus* IT2001 strain on the colonic mucosa of iron-overloaded mice have been reported [7]. Feeding of the *S. thermophilus* YIT2001 strain to mice resulted in high inhibitory activity against lipid peroxidation in liposomes, resulted in a decrease of lipid hydroperoxide in the colonic mucosa [7].

Therefore, lactic acid bacteria that can eliminate environmental lipid hydroperoxide directly, should be vigorously investigated as probiotics to prevent bowel diseases. Previously, we isolated the *Pediococcus pentosaceus* Be1 strain that reduces environmental hydrogen peroxides [8].

Based on this previous isolation method, which was improved upon in this study, we successfully isolated the *Lactobacillus plantarum* P1-2 strain that reduces environmental fatty acid hydroperoxides, which are primary peroxidation products of free fatty acids and are also derived from the hydrolysis of esterified lipid hydroperoxides. We then investigated the effects resulting from the administration of the two isolated lactic acid bacteria strains that reduce environmental hydrogen peroxide (*P. pentosaceus* Be1 strain) and fatty acid hydroperoxide (*L. plantarum* P1-2 strain) in this study. We first examined the inhibitory effects against internal oxidative stress in *C. elegansΔfer-15;mev-1* [9], as the free-living nematode *Caenorhabditis elegans* offers several distinct advantages for aging research at the organismal level [10, 11]. As the distinct inhibitory effects were observed in both strains, we next investigated the effects of the *L. plantarum* P1-2 and *P. pentosaceus* Be1 strains on major organs in mammals, particularly the intestines and colon in an oxidative stress rat model.

## Materials and methods

### Selective isolation medium for lactic acid bacteria

To screen bacteria exhibiting high lipid hydroperoxide-eliminating activity, we used modified GYP medium [8] supplemented with 1% linoleic acid hydroperoxide serving as the fatty acid hydroperoxide. Linoleic acid hydroperoxide was prepared in bulk by oxidizing 250 ml of linoleic acid by incorporating 100% O2 at 70°C. This was suspended in 5% sterilized Tween 80 solution (v/v).

### Identification of lactic acid bacteria

Lactic acid bacteria strains that exhibit high lipid hydroperoxide-eliminating ability were isolated from 86 fermented foods following five cycles of plate culture using the enrichment medium prepared above at 37°C. We identified lactic acid bacteria strains based on taxonomical characteristics such as morphology, fermentation form, catalase, ratio of L-form to D-form in lactic acid production, sugar requirement pattern, and cell wall components [12]. We also identified strains by 16S rDNA sequencing.

### Evaluation of bacterial strains and their culture conditions

*L. plantarum* P1-2, *P. pentosaceus* Be1, other tested strains from fermented foods, and their type strains were cultured under various conditions. Specifically, the *L. plantarum* P1-2 strain, *L. plantarum* NRIC1067^T^, *P. pentosaceus* Be1 strain, *P. pentosaceus* NRIC 0099^T^, *Lactobacillus casei* NRIC 0644^T^, *Lactobacillus delbrueckii* subsp. *bulgaricus* NRIC 1688 ^T^, *Lactobacillus delbrueckii* subsp. *delbrueckii* NRIC 0665 ^T^, *Lactobacillus fermentum* NRIC 1752 ^T^, and *Lactobacillus salivarius* subsp. *salicinius* NRIC 1072 ^T^ were aerobically cultured in GYP medium with shaking at 37°C. *Lactobacillus alimentarius* NRIC 1640^T^, *Lactobacillus ferciminis* NRIC 0492 ^T^, *Lactococcus lactis* subsp. *lactis* NRIC 1149 ^T^, *Leuconostoc mesenteroides* subup. *mesenteroides* NRIC 1541 ^T^, *Weissella viridescens* NRIC 1536 ^T^, *Weissella cibaria* NRI 0527 ^T^, and *P. acidilactici* NRIC 0115^T^ were aerobically cultured in GYP medium with shaking at 30°C. *Lactobacillus acidophilus* NRIC1547^T^ and *S. thermophilus* NRIC0256^T^ were grown in static culture in GYP medium at 37°C. *Lactobacillus brevis* NRIC 1638 ^T^ and *P. damnosus* NRIC 0214 ^T^ were grown in static culture at 30°C in MRS medium. *Bacillus subtilis* NRIC 1015 and *Escherichia coli* NRIC 1509 were grown aerobically in NB medium at 37°C.

### Evaluation of alkyl and fatty acid hydroperoxide-eliminating activity by lactic acid bacteria

Bacterial cells were harvested at their late logarithmic or early stationary growth phase by centrifugation and washed with 50 mM sodium phosphate buffer (pH 7.0). The late logarithmic or early stationary growth phase was determined based on optical density at 660 nm. After the value of the cell suspension was adjusted to 1.6, it was used for determining the dry cell weight and measuring the hydroperoxide-eliminating activity. For determination of the dry cell weight, 200 ml of the bacterial cell suspension was centrifuged at 48,400 × *g* for 10 min, and the cell pellet was dried at 100°C until a constant weight was achieved (S1 Table).

To measure hydroperoxide-eliminating activity, the remaining bacterial suspension was incubated with 0.3, 1.0, or 3.0 mM cumene hydroperoxide serving as alkyl hydroperoxides, or 0.25, 0.5, or 1.0 mM fatty acid hydroperoxide serving as linoleic acid hydroperoxide in the presence of 50 mM glucose for 1.5 h at 37°C with shaking. After the reaction was terminated by centrifugation to remove the bacterial cells at 4°C, remaining hydroperoxides were identified by the method described below.

The fatty acid hydroperoxide used in this assay was prepared based on the method of lipoxygenase oxidation of linoleic acid [13] (Funk *et al*., 1976). After incorporation with 100% O2, the linoleic acid mixture was extracted using diethylether, which was then removed by evaporation. To evaluate linoleic acid hydroperoxide-eliminating activity, linoleic acid hydroperoxide was dissolved in a 2.5% Triton-XP-100 solution.

### Determining the concentration of identified cumene hydroperoxide and linoleic acid hydroperoxide

To determine the remaining cumene hydroperoxide or fatty acid hydroperoxide concentration after the reaction with living cells, we applied the modified ferric thiocyanate assay [14]. The ferric thiocyanate mixture consisted of 960 µl chloroform:methanol (2:1 v/v), 40 µl of supernatant containing cumene or lipid hydroperoxide, and 200 µl colorimetric reaction mixture. The colorimetric reaction mixture contained 3% KSCN/methanol and 4.5 mM FeSO4·7H2O/0.2 N HCl (3:1 v/v). Each assay mixture was added at 25°C for the colorimetric reaction. After 5 min, the solution was removed by centrifugation, followed by a spectrophotometric measurement at 500 nm. A calibration curve with cumene hydroperoxide was generated when we evaluated the cumene hydroperoxide-eliminating activity. Calibration curves for 13S-hydroperoxy-9z and 11E-octadecatienoic acid (13-HpODE, Cayman Chemical) were also prepared to evaluate linoleic acid hydroperoxide-eliminating activity.

We also analyzed the reaction product of cumene hydroperoxide by HPLC. An HPLC system equipped with a PEGASIL ODS C-18 (4.6 mm × 250 mm, Senshu Scientific co., ltd.) reverse phase HPLC column, L-7100 pump (Hitachi), L-7420 UV-VIS detector (Hitachi), and D-7500 recorder (Hitachi) was used, and the injection volume was 100 µl. The products were eluted with acetonitrile:5 mM potassium phosphate buffer, pH 7 (3:7), at a flow rate of 1 ml/min, at 40°C with monitoring at 265 nm. Product elution peaks were identified by comparing authentic standards under identical elution conditions.

Additionally, we analyzed the reaction product of linoleic acid hydroperoxide by HPLC. The HPLC system was equipped with a Jupiter 5 µm C18 (300 Å 250 mm × 4.6 mm, Phenomenex) reverse phase HPLC column, LC-20A pump (Shimadzu), SPD-20A PDA detector, and CBM-20 controller. The injection volume was 5 µl. The products were eluted with 1 g/L acetic acid:acetonitrile:tetrahydrofuran (52:30:18) at a flow rate of 0.8 ml/min at 40°C with monitoring at 234 nm. Product elution peaks were identified by comparing authentic standards, specifically 13-HpODE (13S-hydroperoxy-9Z, 11E-octadecadienoic acid; Cayman Chemical) and 13-HODE (13S-hydroxy-9Z, 11E-octadecadienocic acid; Cayman Chemical), under identical elution conditions.

### Animal test 1: Evaluating the lifespan of the short-lived, oxygen-sensitive *C. elegans* mutant

In this study, we evaluated the lifespan of *C. elegans* with mutations in both *fer-15* and *mev-1*. The *C. elegans fer-15* mutant was sterile when grown at 25°C, as under these conditions spermatids failed to activate into spermatozoa. Mutations in *mev-1* render animals hypersensitive to high oxygen concentrations due to increased superoxide levels [15]. These mutant *C. elegans* also accumulate more fluorescent material (lipofuscin) with age [16]. The *C. elegans Δfer-15;mev-1* strain was obtained from the Tokai University School of Medicine Basic Medical Science and the Molecular Medicine Department of Molecular Life Sciences. We administered lactic acid bacteria strains that have high or low hydroperoxide-eliminating activity to *C. elegans Δfer-15;mev-1*, a low lifespan mutant with high oxygen sensitivity.

We defined four administration groups of tested bacteria strains. These included the *E. coli* OP50 strain as the control group (OP50 group), the *L. plantarum* P1-2 strain that demonstrates high fatty acid hydroperoxide-reducing ability (P1-2 group), the *P. pentosaceus* Be1 strain that has high hydrogen peroxide-reducing ability (Be1 group), and the *S. thermophilus* NRIC0256^T^ strain that exhibits low hydroperoxide eliminating ability (ST group). Animals were cultured on nematode growth medium NGM agar plates seeded with the *E. coli* OP50 strain at 20°C. Embryos (eggs) were collected from young adult hermaphrodites on NGM agar plates using alkaline sodium hypochlorite [17]. The released eggs were allowed to hatch through overnight incubation at 20°C in S buffer [18]. We continuously grew young stage nematodes until the L4 stage on NGM agar plates (90 mm), with live bacteria (*Escherichia coli* strain OP50) added as food. L4 stage nematodes were transferred to 10 modified GYP medium agar plates (30 mm) that contained 100 mM MES at pH 6.0, with each live bacteria group (*E. coli* OP50 strain, *L. plantarum* P1-2 strain, *P. pentosaceus* Be1 strain, and *S. thermophilus* NRIC0256^T^) and the lifespan at 25°C was evaluated to prevent progeny production. Death was defined as the loss of spontaneous movement and lack of response to touch with a platinum wire.

Statistical analysis was carried out by Student’s *t*-test and Tukey’s multiple-range test. The least significant difference test was used for means separation at *P* < 0.05 within strains.

### Animal test 2-1: Administration of lactic acid bacteria to iron-overloaded rats experiencing induced lipid peroxidation

All animal experiments were performed with permission from the Committee on Animal Experiments of Tokushima University (permit number: 11,013) according to the guidelines for the care and use of laboratory animals set by the University (Tokushima, Japan).

Wistar rats (6-week-old male, Japan SLC, Shizuoka, Japan) were maintained in a room at 23 ± 1°C on a 12-h light–dark cycle. Rats were maintained on AIN-76 as their basal diet. In the experimental phase, we administered various diets. Specifically, the control group received the basal diet and 5% skim milk powder, the Fe group received the basal diet plus 0.5% ferrous fumarate and 5% skim milk powder, and the LAB group received basal diet plus 0.5% ferrous fumarate and 5% lyophilized lactic acid bacteria powder (S2 Table). Lyophilized lactic acid bacteria powder consists of a 1:9 ratio of dried lactic acid bacteria cells:skimmed milk powder at approximately 10 × 10^9^ cfu/g. This powder was mixed with the basal diet and stored at −18°C until the experimental phase.

The rats had free access to food and water, and the food was replaced every 24 hours. After one week of AIN-76 diet treatment, two weeks of iron-enriched diets including lactic acid bacteria were administered. Before the experimental phase, rats received AIN-76 for one week. In the experimental phase, rats maintained the control group diet, Fe group diet, or the LAB group diet for two weeks. Rats were randomly assigned to each group.

Body weight was recorded daily, and after the dietary treatments, the rats were anesthetized using diethyl ether. Rats were sacrificed by cardiocentesis, and blood was collected with heparin sodium on ice and then centrifuged. The abdomens were opened along the median line, and the stomach, intestines, colon, and liver were rapidly excised and rinsed gently with ice-cold saline. Stomach, intestines, and colon were opened longitudinally to collect the respective mucosa.

### Animal test 2-2: Determination of malondialdehyde in rat organs

The stomach, intestines, colonic mucosa, and liver were prepared as homogenates on ice. Each homogenate was determined by malondialdehyde [19], and the total protein concentration was quantified using the Bradford method. We represented lipid peroxidation level as MDA/mg of protein. Data are expressed as the mean ± SD. Differences between the control and iron fumarate group were analyzed by unpaired *t*-test. Data obtained from over three groups were analyzed using non-repeated analysis of variance (non-repeated ANOVA). When the result of non-repeated ANOVA was significant (*P* < 0.05), Student–Newman–Keuls methods were conducted (*P* < 0.05).

### Nucleotide sequence accession number

The 16S rDNA sequence of the *L. plantarum* P1-2 strain was submitted to the DNA Data Bank of Japan under accession number LC0424332.

## Results

### Isolation of lactic acid bacteria eliminating alkyl and fatty acid hydroperoxide and distribution of their eliminating activity

Although the enrichment medium contained sources of ROS, 116 strains of the isolates grew well and were isolated from various kinds of fermented foods (Table 1). The obtained isolates included 75 strains of *lactobacilli*, 24 strains of *Pediococci*, and 17 strains of *Leuconostocs*. Next, we measured the eliminating activity of cumene hydroperoxide and linoleic acid hydroperoxide in the 116 isolates. We successfully isolated one strain from the leaven, and this strain displayed the highest eliminating activity for both substrates (Fig 1A and 1B). Based on the taxonomical characterization [12] and the 16S rDNA sequence of this strain, we identified it as *L. plantarum* P1-2 (S3 Table).

**Table 1.** Isolation of lactic acid bacteria exhibiting high eliminating activity for environmental hydroperoxides from fermented foods.

|  | Sources for screening |  |  |  |  |  |  | Total number |
| --- | --- | --- | --- | --- | --- | --- | --- | --- |
|  | Rice malts | Rice bran | Fermented vegetables with salts | Salted radish aged with koji rice | Leaven | Seafood | Others |  |
| Number of sources for screening | 26 | 28 | 13 | 4 | 8 | 4 | 3 | 86 |
| Isolates in 1st isolation step | 352 | 198 | 38 | 183 | 282 | 15 | 7 | 1075 |
| Isolates after 5th isolating step with enrichment | 80 | 11 | 1 | 8 | 30 | 15 | 3 | 148 |
| Isolates selected by cumene hydroperoxide and linoleic acid hydroperoxide eliminating activity | 29 | 25 | 10 | 6 | 24 | 15 | 7 | 116 |

**Fig 1.**
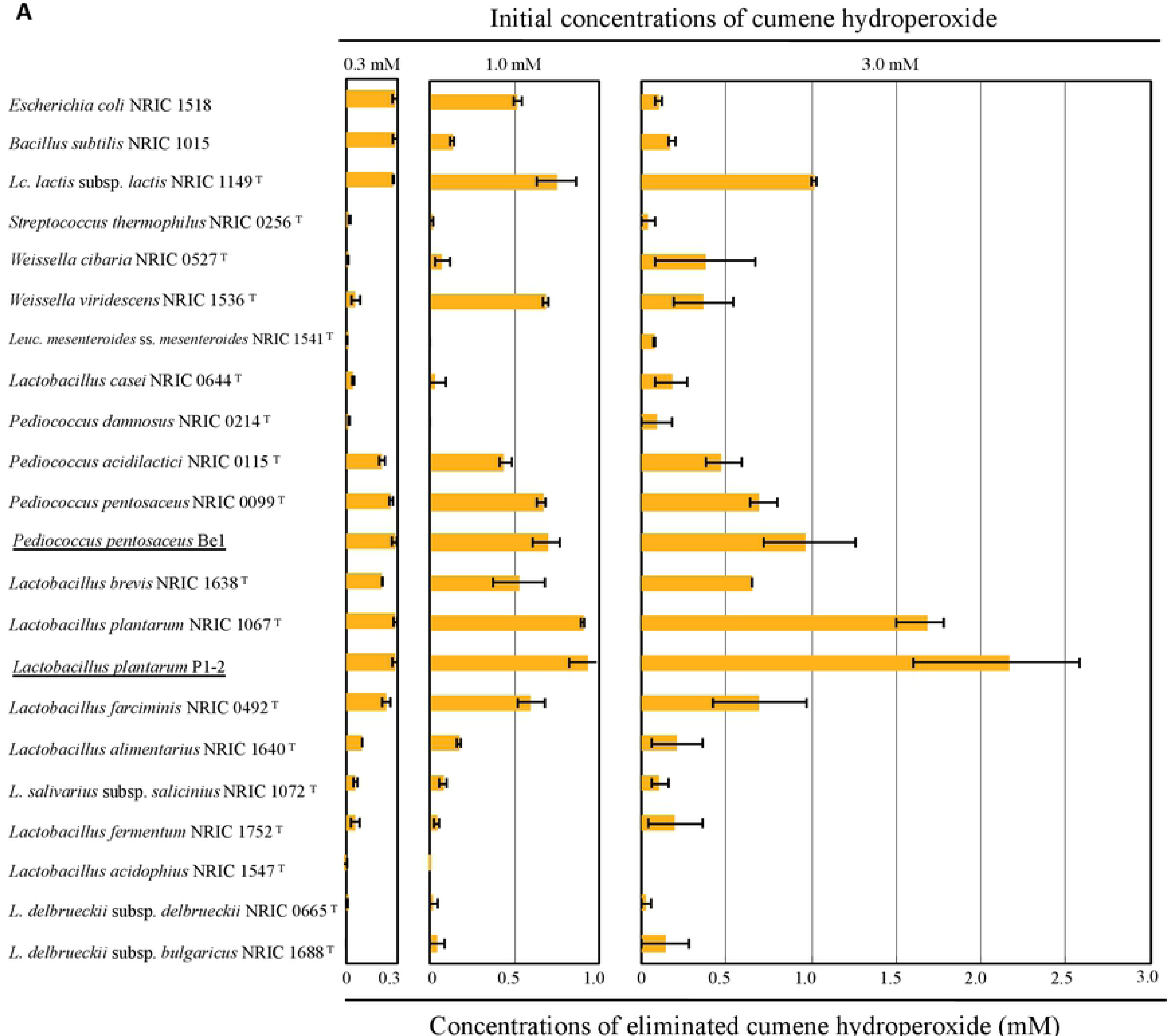
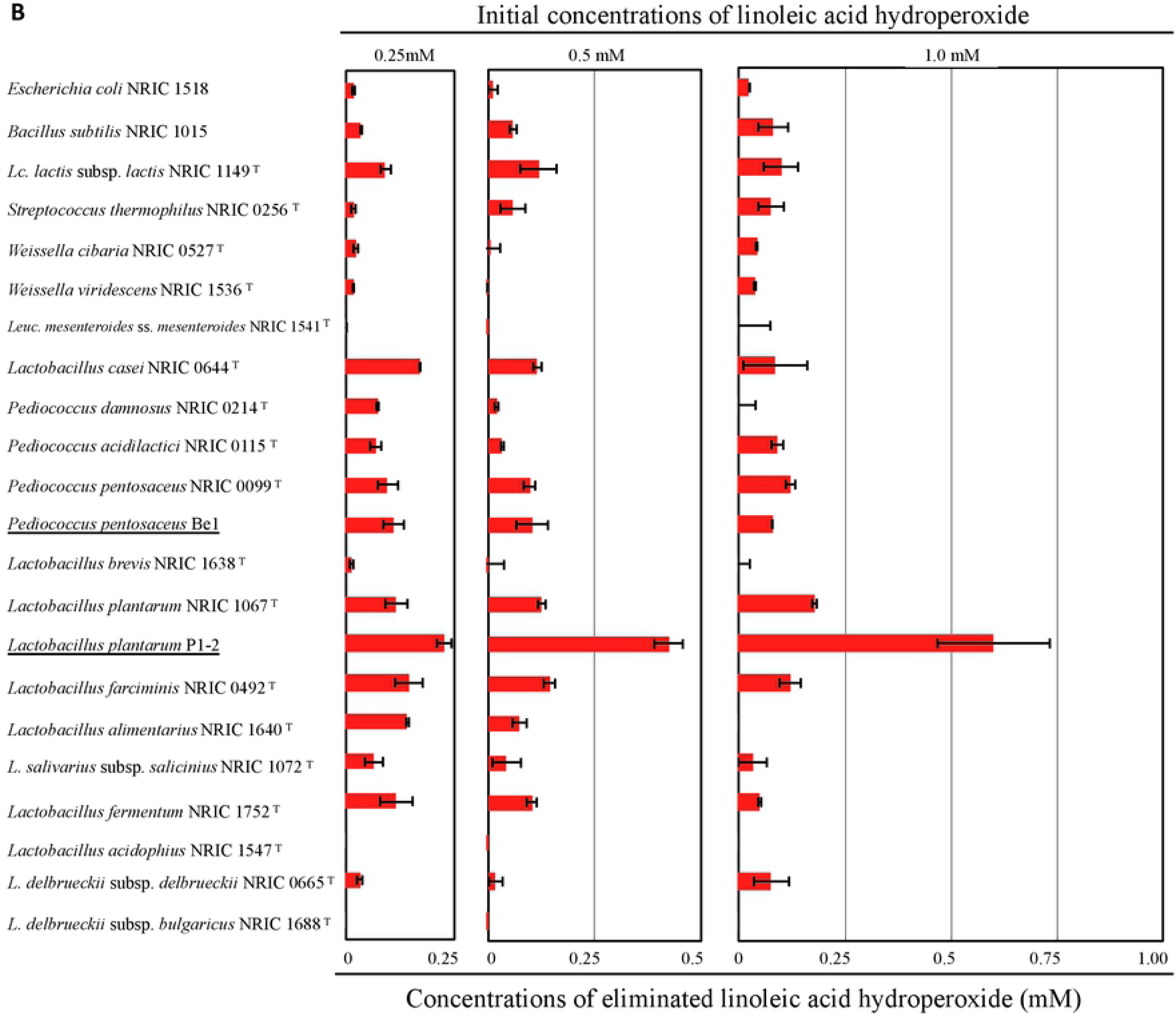
The distribution of eliminating activities for cumene and linoleic hydroperoxide in lactic acid bacteria and the difference in their capacities. (A) Twenty lactic acid bacterial strains and typical bacterial species were cultured under specific conditions described in the Materials and Methodssection. Eachlivingcellwasexposedtovariousconcentrationsofcumene hydroperoxide at 0.3, 1.0, and 3.0 mM for 1.5 h. After treatment, the extent of decomposition of the hydroperoxide was calculated as their eliminating activity by the difference of the initial concentration and the remaining concentration in the culture medium. The bar graph represents the mean values from three independent experiments, and error bars indicate the standard deviation (SD). (B) The same twenty lactic acid bacterial strains and typical bacterial species as (A) were cultured under specific conditions described in the Materials and Methods section. Each living cell was exposed to various concentrations of linoleic acid hydroperoxide at 0.25, 0.5, and 1.0 mM for 1.5 h at 37°C. After treatment, the extent of decomposition of the hydroperoxide was calculated as (A).

We compared hydroperoxide eliminating capacities of the *L. plantarum* P1-2 strain, *P. pentosaceus* Be1, and typical strains of lactic acid bacteria applied to the food industry, including fermentation. Using cumene hydroperoxide, the eliminating activities for the substrate were widely preserved in *L. plantarum, P. pentosaceus*, including *P. pentosaceus* Be1 strain, and *L. lactis* (Fig 1A). However, high eliminating activity for linoleic acid hydroperoxide was specifically detected in *L. plantarum*. Specifically, the *L. plantarum* P1-2 strain eliminated over 0.5 mM out of 1.0 mM linoleic acid hydroperoxide in 1.5 h (Fig 1B). Both activities were not detected in dead cells after heat treatment at 100°C for 10 min (Data not shown).

### The relationship between the tolerance and eliminating activity of alkyl hydroperoxide and linoleic acid hydroperoxide-eliminating activity in lactic acid bacteria

To investigate the relationship between the tolerance and eliminating activity of alkyl hydroperoxide and lipid hydroperoxide-eliminating activity in lactic acid bacteria, we compared the number of living cells and the reducing activities for cumene and linoleic acid hydroperoxide per dry cell weight (Fig 2A and B). The cells were treated with various concentrations of hydroperoxides.

**Fig 2.**
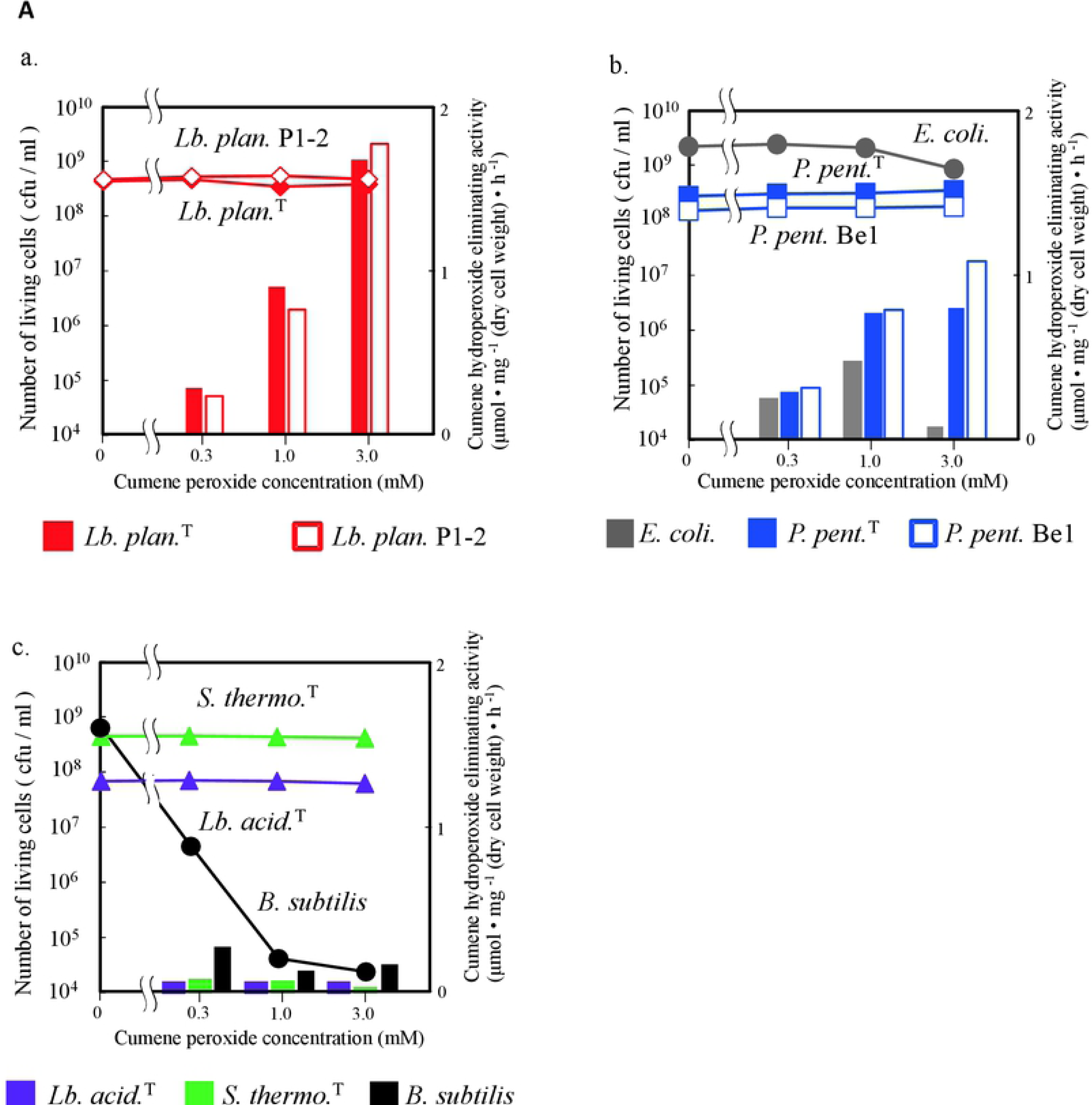
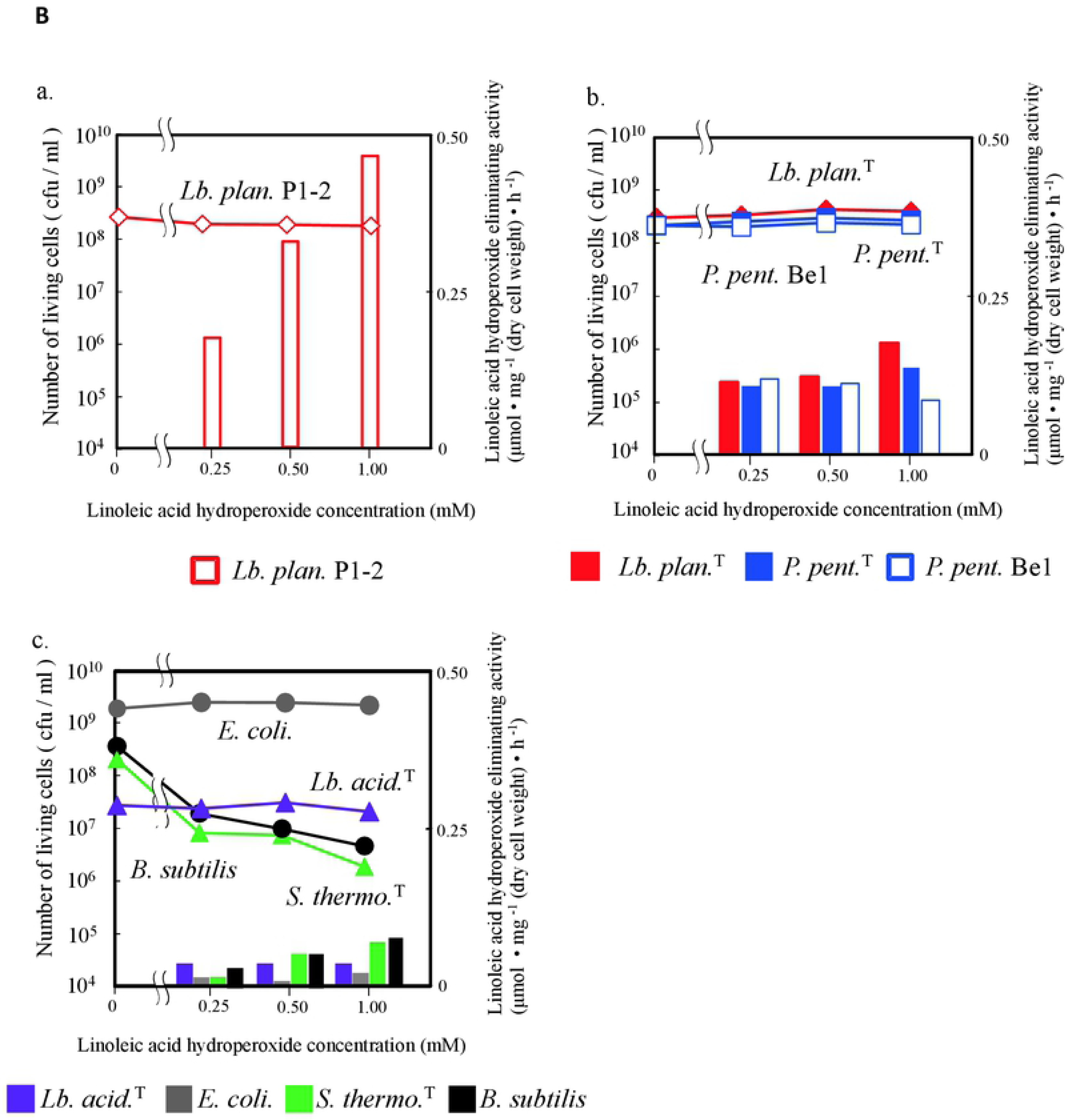
Relationship between the eliminating activity and survival rate of lactic acid bacteria in the presence of cumene or linoleic acid hydroperoxide. (A) Bacterial cells were incubated with various concentrations of cumene hydroperoxide from 0.3 to 3 mM for 1.5 h. After treatment, the cells were diluted and plated, and then the developed colonies were defined as living cells. The eliminating activity for substrate was estimated as described in Figure 1. The relationship between the number of living cells (lines) and the eliminating activity for hydroperoxide (bars) were plotted in each case. The data are mean values of three independent experiments. (a) *L. plantarum* NRIC 1067^T^ and *L. plantarum* P1-2 strain (b) *E. coli* NRIC1518, *P. pentosaceus* NRIC 0099^T^, and *P. pentosaceus* Be1 strain, (c) *L. acidophilus* NRIC 1547^T^, *S. thermophilus* NRIC 0256^T^ and *B. subtilis* NRIC 1015. (B) The same bacterial cells as (A) were incubated with various concentrations of linoleic acid hydroperoxide from 0.25 to 1 mM for 1.5 h at 37°C. After treatment, the cells were diluted and plated, and then the developed colonies were defined as living cells. The eliminating activity for substrate was estimated as described in (A). (a) *L. plantarum* NRIC 1067^T^ and *L. plantarum* P1-2 strain (b) *E. coli* NRIC1518, *P. pentosaceus* NRIC 0099^T^, and *P. pentosaceus* Be1 strain, (c) *L. acidophilus* NRIC 1547^T^, *S. thermophilus* NRIC 0256^T^ and *B. subtilis* NRIC 1015.

The *L. plantarum* NRIC1067^T^ and *L. plantarum* P1-2 strains sustained both the alkyl hydroperoxide reducing activity and the number of living cells in the presence of high concentrations of cumene hydroperoxide of up to 3.0 mM (Fig 2Aa). *E. coli* NRIC1519, *P. pentosaceus* NRIC 0099^T^, and *P. pentosaceus* Be1 strains also exhibited the same tolerance capacity as *L. plantarum* NRIC1067^T^ and *L. plantarum* P1-2 under 3.0 mM cumene hydroperoxide. The eliminating activity for cumene hydroperoxide, however, was much lower than that of *L. plantarum* NRIC1067^T^ and *L. plantarum* P1-2 (Fig 2Ab). Although *S. thermophilus* NRIC0256^T^ and *L. acidophilus* NRIC1547^T^ also retained their cell viability with each cumene hydroperoxide concentration, the number of *B. subtilis* NRIC1015 viable cells decreased under the same conditions. These strains showed low cumene hydroperoxide eliminating activity (Fig 2Ac).

For linoleic acid hydroperoxide, the *L. plantarum* P1-2 strain exhibited potent eliminating activity in the presence of a high concentration of linoleic acid hydroperoxide, up to 1.0 mM while retaining the number of living cells (Fig 2Ba). Similar behaviors were also observed in *P. pentosaceus* NRIC 0099^T^ and *P. pentosaceus* Be1 strains. However, the eliminating activities nearly plateaued at 0.25 mM linoleic acid hydroperoxide and were much lower than those of the *L. plantarum* P1-2 strain (Fig 2Bb). *E. coli* NRIC1519 also retained a significant number of living cells against each linoleic acid hydroperoxide. In contrast, the numbers of *S. thermophilus* NRIC0256^T^, *L. acidophilus* NRIC1547^T^, and *B. subtilis* NRIC1015 decreased with increased concentration of linoleic acid hydroperoxide from 0.25 to 3.0 mM. These strains exhibited generally low linoleic acid hydroperoxide eliminating activity (Fig 2Bc).

The *L. plantarum* P1-2 strain clearly showed higher reducing activity for both cumene and linoleic acid hydroperoxide than did other strains. Also, we examined both hydroperoxide reducing reaction products by HPLC. The *L. plantarum* P1-2 strain converted cumene hydroperoxide to 2-phenyl-2-propanol (Fig 3A) and reduced 13-HpODE to 13-HODE (Fig 3B). These results indicate that the *L. plantarum* P1-2 strain reduces cumene and linoleic acid hydroperoxide by a two-electron reduction.

**Fig 3.**
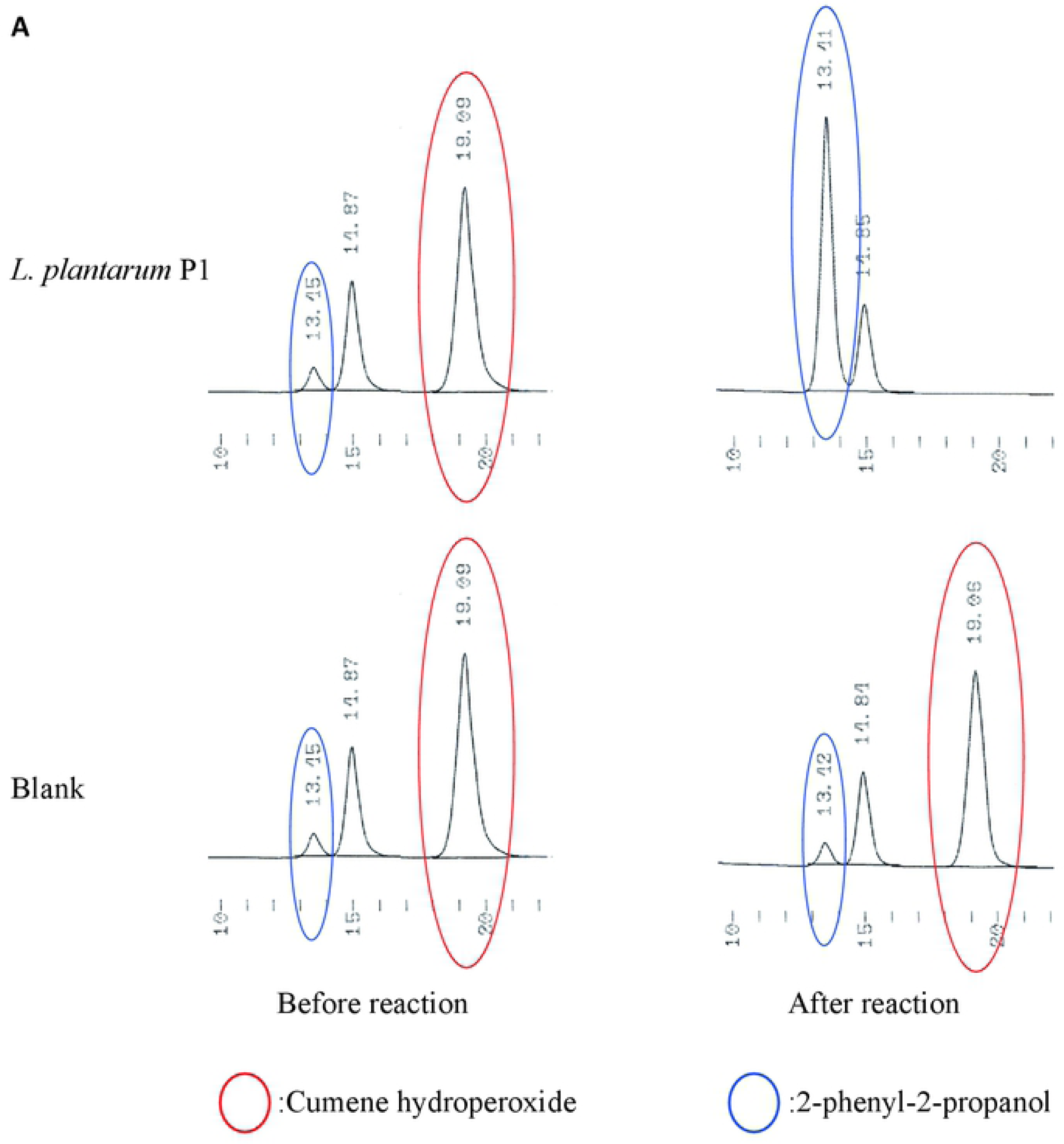
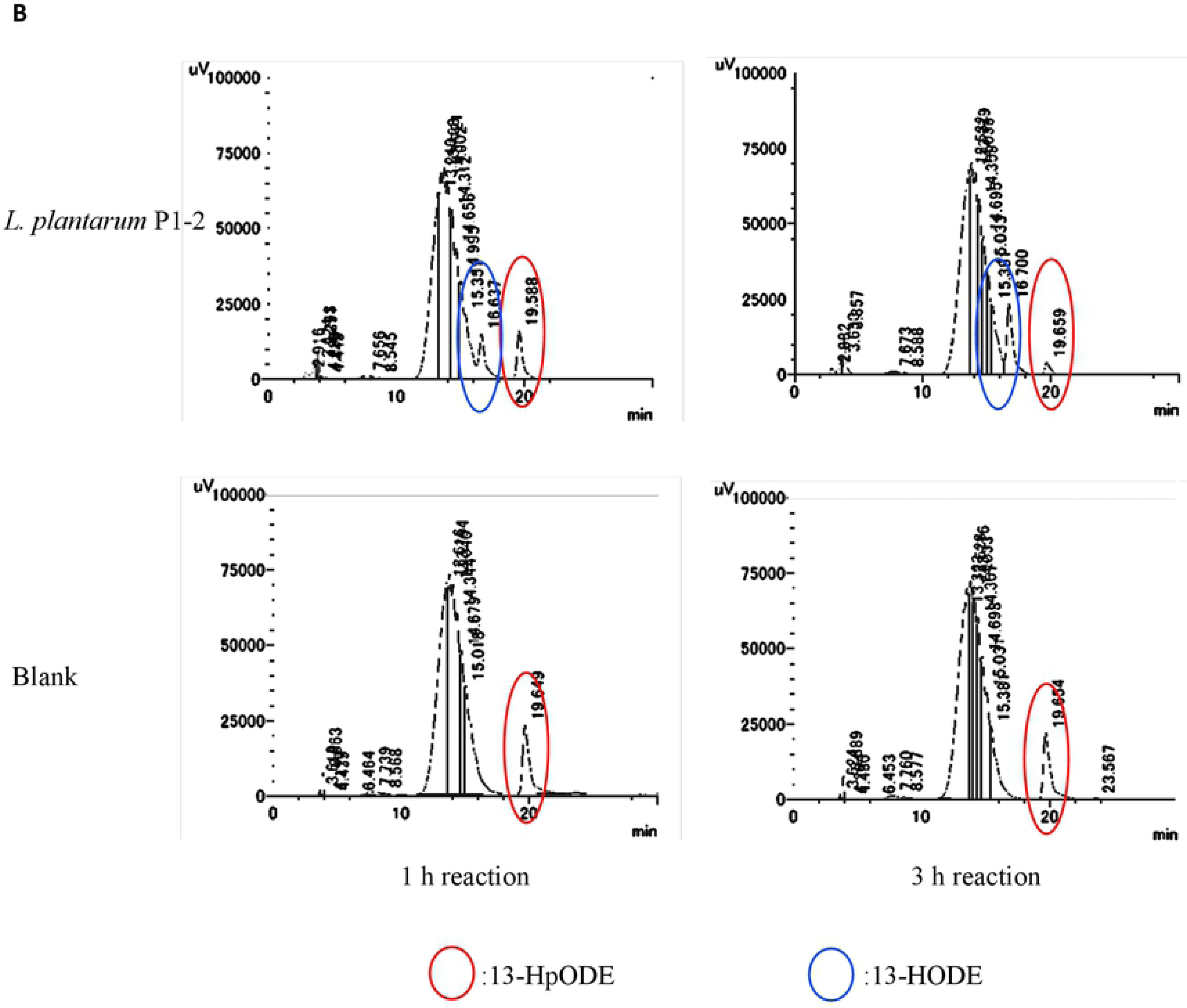
*Lactobacillus plantarum* P1-2 reduces cumene and linoleic acid hydroperoxide to the corresponding hydroxyls. (A) *L. plantarum* P1-2 was aerobically cultured in GYP medium, and the cells were incubated in 50 mM sodium phosphate (pH 7) containing 25 mM glucose and 3 mM cumene hydroperoxide for 1.5 h at 37°C. After the reaction, each metabolite was analyzed by HPLC equipped with an ODS column. Cumene hydroperoxide and the corresponding alcohol (2-phenyl-2-propanol) were eluted at retention times of 19 and 13 min, respectively. (B) *L. plantarum* P1-2 was aerobically cultured in GYP medium, and the cells were incubated in 50 mM sodium phosphate (pH 7) containing 25 mM glucose 3 mM 13-HpODE (2) for 3 h at 37°C. 13-HpODE was used as the stable substrate of linoleic acid hydroperoxide. After the reaction, each metabolite was analyzed by HPLC equipped with an ODS column. 13-HpODE and the corresponding hydroxyl (13-HODE) were eluted with retention times of 19 and 16 min, respectively.

### Animal test 1: Evaluating the lifespan of the short-lived, oxygen-sensitive *C. elegans* mutant

To investigate the efficacy of the reductase activity for fatty acid hydroperoxide activity of *Lactobacillus plantarum* P1-2 *in vivo*, survival analyses using nematode (*Caenorhabditis elegans*) were performed. *L plantarum* P1-2 was administered to *C. elegans Δfer-15* at L4 stage, and its life span was monitored. *E. coli* OP50, *S. thermophilus* NRIC0256^T^ and *P. pentosaceus* Be1 were also investigated. This assay was conducted on GYP medium instead of NGM medium, as all tested bacteria grow well and exhibit high fatty acid hydroperoxide activity on GYP compared to that observed on NGM. Additionally, the pH of GYP was fixed at pH 6 with MES buffer to allow nematodes to live together with the bacteria and assimilate with them without the effects of bacterial metabolites such as organic acids. Under this condition, *L. plantarum* P1-2 and *P. pentosaceus* Be1 were likely to prolong the life span of *C. elegans Δfer-15* more significantly than were *S. thermophilus* NRIC0256^T^ and *E. coli* OP50, respectively (S1 Fig). To clarify if this life extension is associated with fatty acid hydroperoxide or hydrogen peroxide reductase activity, the four bacterial strains were administered to a *C. elegans Δmev1* mutant that exhibits oxygen-sensitivity. As a result, *L. plantarum* P1-2 and *P. pentosaceus* Be1 significantly extended the life span of the mutant compared to *S. thermophilus* NRIC0256^T^ (p<0.001) (Fig 4), consistent with the capacity to reduce hydroperoxide *in vitro* (Fig 1). This result suggested that *L. plantarum* P1-2 and *P. pentosaceus* Be1 are effective for reducing oxidative stresses *in vivo*. To determine the specific organs involved in these bacterial administrations, further experiments were performed.

**Fig 4.**
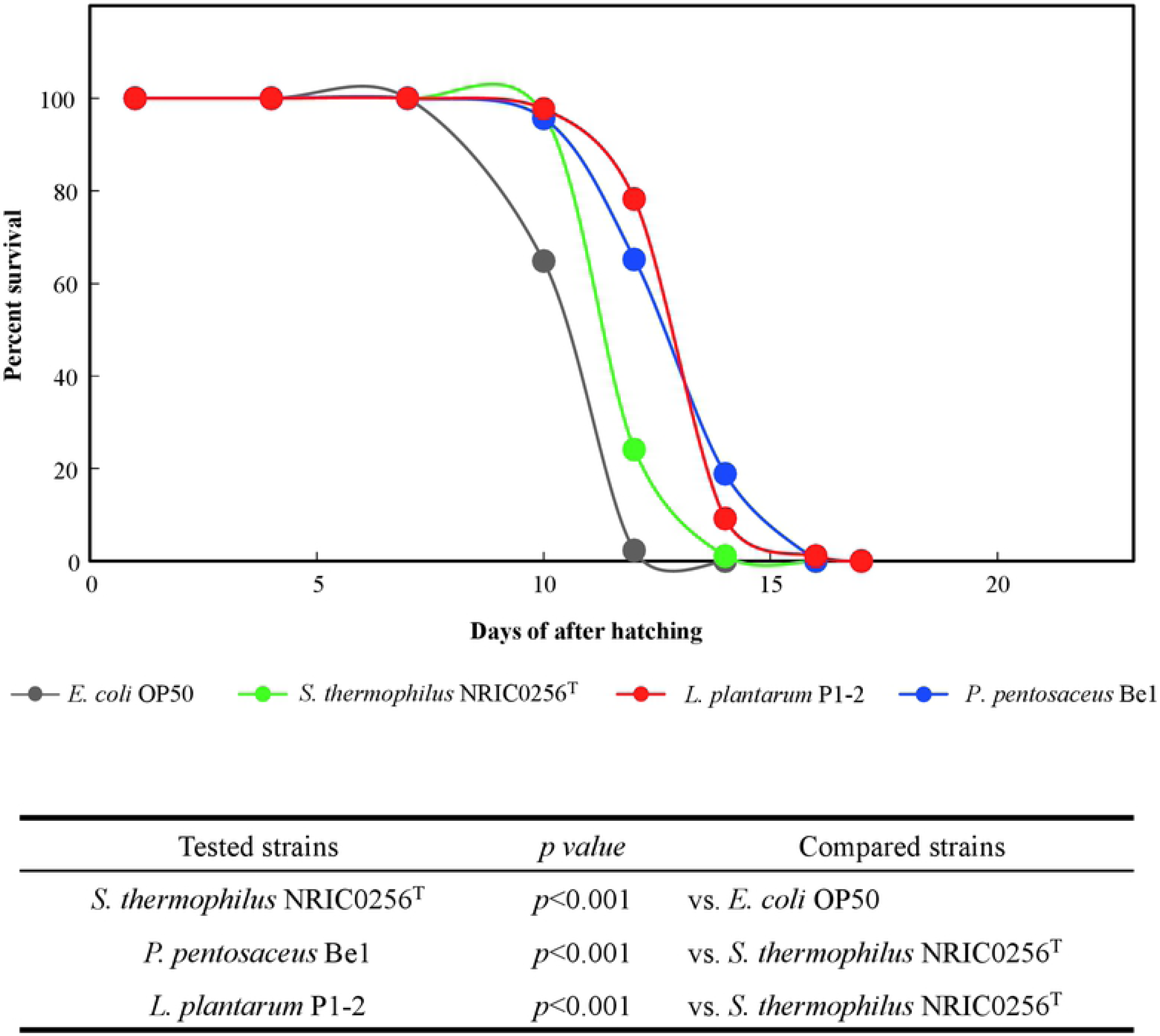
Prolongation of the lifespan of *C. elegans Δfer-15;mev-1* with lactic acid bacteria. *E. coli* OP50 (gray), *S. thermophiles* NRIC0256T(green), *L. plantarum* P1-2 (red), and *P. pentosaceus* Be1 (blue) were administered to *C. elegans Δfer-15;mev-1* at the growth stage L4. The mutants were hatched on pH stat GYPmedium, andtheir lifespanwasmonitoreduntilannihilation. Statistical analysiswascarriedoutby Student’s t-test and Tukey’s multiple-range test. The least significant difference test was used for means separation at *P* < 0.05 within each strain. One hundred animals were measured for each strain at 25°C.

### Animal test 2: The effect of lactic acid bacteria administration on the eliminating activity of hydroperoxides in iron-loaded rats using the colonic mucosal lipid peroxidation model

Some studies have shown that iron administration increases the levels of lipid peroxidation markers in rat liver [20]. Similarly, iron increases the levels of lipid a peroxidation marker in the colonic mucosa of mice [7]. Iron induces the production of reactive oxygen species (ROS), followed by ROS-induced gastrointestinal mucosa lipid peroxidation [21] and oxidative stress-induced tissue damage [22]. When we examined the effects of 0.2% and 0.5% iron fumarate in Wister rats (n = 3), colonic mucosa and liver homogenate MDA increased in a dose-dependent manner. We settled on an iron fumarate concentration for lipid peroxidation of 0.5%. We also evaluated the increase of MDA in the gastric mucosa, intestinal mucosa, colonic mucosa, and liver homogenates and serum in iron fumarate overloaded rats (n = 9). For the gastric mucosa, intestinal mucosa homogenate, and serum, there was no significant increase of MDA. Conversely, colonic mucosa and liver homogenate both exhibited a significant increase in MDA (Fig 5). These results suggest that iron overload in rats increases lipid peroxidation in the liver and colonic mucosa.

**Fig 5.**
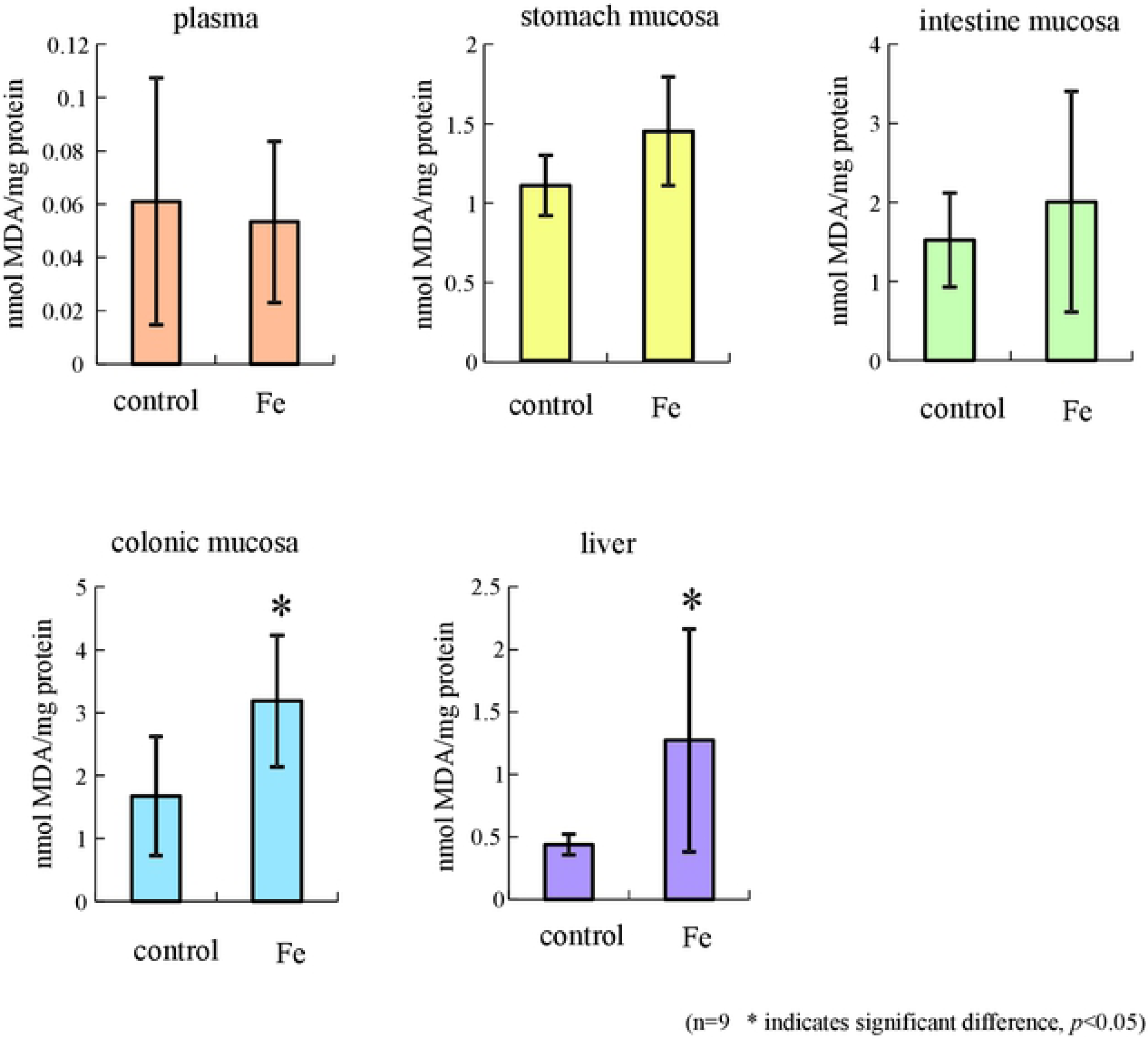
Lipid peroxidation levels of plasma and tissues obtained from iron-overloaded rats. Lipid peroxidation in rats was induced by 0.5% iron fumarate (Fe), and the MDA level was measured in each organ. Data are expressed as the mean values ± SD (n = 9). The asterisk (*) indicates a statistical difference of *P* < 0.05.

We also compared the effect of *P. pentosaceus* Be1 group, *L. plantarum* P1-2 group, and *S. thermophilus* NRIC0256^T^ group on iron overloaded rats (n = 4). MDA levels were lower in the *L. plantarum* P1-2 group than in the *S. thermophilus* NRIC0256^T^, and *P. pentosaceus* Be1 groups (S2 Fig). Next, we compared confirmatively the effect of *L. plantarum* P1-2 and *S. thermophilus* NRIC0256^T^ on iron overloaded rats (n = 6). The MDA levels following administration of *L. plantarum* P1-2 were significantly lower in the colonic mucosa than those of *S. thermophilus* NRIC0256^T^ (Fig 6). We examined the administration effect of heat-treated dead *L. plantarum* P1-2 and *S. thermophilus* NRIC0256^T^ cells on iron overloaded rats. Deadlactic acid bacteria cells exert no significant effect on production of MDA (data not shown). These results indicate that administration of living *L. plantarum* P1-2 cells decreases lipid peroxidation in the colonic mucosa.

**Fig 6.**
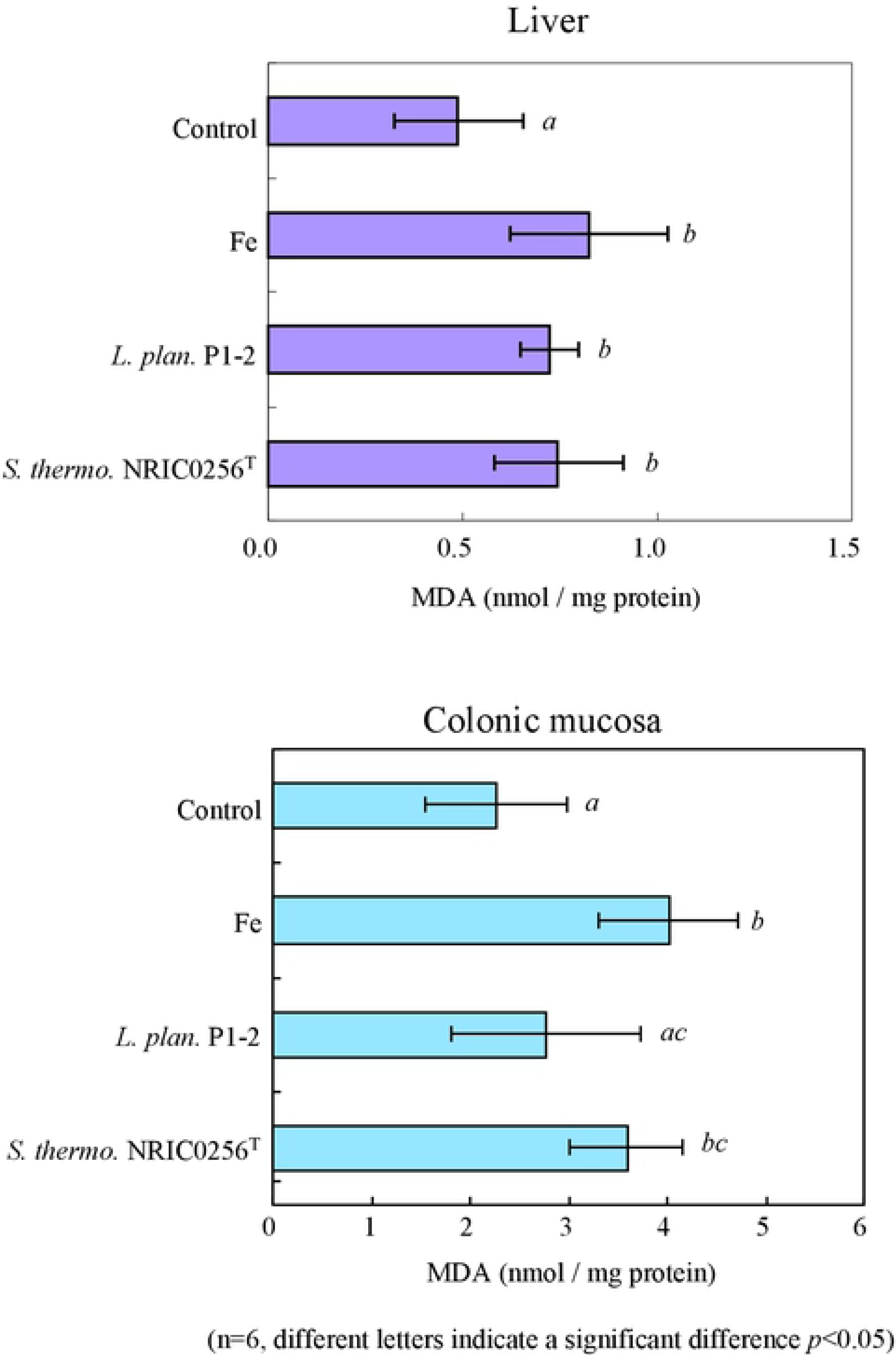
The effect of *Lactobacillus plantarum* P1-2 administration on the liver and colonic mucosa of iron-overloaded rats. *L. plantarum* P1-2 was administered to iron-overloaded rats, and the MDA levels in the liver and colonic mucosa were compared to those of the healthy (control), iron-overloaded rats (Fe). *S. thermophilus* NRIC0256^T^ was also tested as the control strain. The data are the mean values ± SD (n = 6), and the different letters indicate the statistical significance at *P* < 0.05.

## Discussion

Hydroperoxides, such as hydrogen peroxide and lipid hydroperoxides, are toxic and thought to contribute to various diseases and the aging process. Two isolated lactic acid bacteria, *P. pentosaceus* Be1[8] and *Lactobacillus plantarum* P1-2, show a 2-electron reduction of hydrogen peroxide to water and of fatty acid hydroperoxides, a type of lipid hydroperoxide, to the corresponding hydroxy derivatives. The oxygen-sensitive and short-lived nematode mutant, *C. elegans Δfer-15;mev-1*, accelerates the accumulation of hydroperoxides inside the body due to a lack of superoxide dismutases. The administration of *P. pentosaceus* Be1 and *L. plantarum* P1-2 should decrease hydroperoxides *in vivo*, and with both strains we observed this and an extension of nematode lifespan.

To elucidate which organs are affected by these bacterial administrations, the TBARS assay for evaluating the oxidative stress tolerance by rat organs was established. Based on the assay condition used in mice [23], we defined the proper dose of iron(II)-fumarate in rats, and we measured the MDA levels in each organ. Our results demonstrated significant increases of MDA levels in the colonic mucosa and liver, a finding that is consistent with previous reports on colonic mucosa in mice [23] and liver in rat [20, 24]. In contrast, the increase of the MDA level in stomach and intestine mucosa of rats was not observed. This is likely because in the stomach, iron(II)-fumarate was easily dissolved in gastric juices and then rapidly diffused into the intestine. Alternatively, lipid hydroperoxides may have been metabolized in the stomach prior to their accumulation in the tissue cells, as the turnover of intestine mucosa cells is much higher than that in colonic mucosa. Therefore, our findings are in agreement with the idea that the above specific lipid peroxidation in colonic mucosa is likely due to its vulnerability to oxidative stress compared to that of other gastrointestinal mucosa [1].

The administration of *L. plantarum* P1-2 to iron-overloaded rats resulted in a significant decrease in MDA levels in the colonic mucosa, and administration of *P*. *pentosaceus* Be1 did not cause this effect. The accumulation of MDA by a ferrous iron is thought to be primarily responsible for the stimulation of the Fenton reaction and subsequent accumulation of lipid hydroperoxides. Thus, it is suggested that *L. plantarum* P1-2 possesses the ability to inhibit the Fenton reaction (related reactive oxygen) and/or to stimulate the reduction activity for lipid hydroperoxides (related free radical). Ito and co-workers also discussed the decrease of MDA levels in iron-overloaded mice by the administration of *S. thermophiles* YIT2001 as being related to the potential to eliminate reactive oxygen or free radicals [7].

In this study, the decrease of MDA levels following bacterial administration was observed in the case of *L. plantarum* P1-2 exhibiting high reductase activity for exogenous fatty acid hydroperoxides but was not observed in *P. pentosaceus* Be1 possessing low activity for these molecules. It is established that a large amount of unsaturated lipids such as linoleic acids are contained in the digestive tract of humans, where lipid peroxidation is evoked by endogenous or exogenous reactive oxygen species [1]. Thus, this result suggests that the reductase activity of *L. plantarum* P1-2 for exogenous fatty acid hydroperoxides is effective for decreasing MDA *in vivo*. This idea is also supported by the observation that the accumulation of lipofscin derived from lipid hydroperoxides in *C. elegans Δfer-15;mev-1* was inhibited by administrating *L. plantarum* P1-2 more effectively than by administering *P. pentosaceus* Be1 (data not shown).

Although there are several excellent antioxidant investigations in lactic acid bacteria [25-30], to our knowledge, a report describing the reduction of exogenous alkyl hydroperoxides and linoleic acid hydroperoxides by bacteria has not been published. In this study, we demonstrated that bacteria exhibit reduction ability against both hydroperoxides; however, it is unclear if *L. plantarum* P1-2 can directly reduce esterified fatty acids such as phospholipid hydroperoxide and cholesterol. The molecular mechanism by which bacteria reduce hydroperoxides requires further investigation. Based on these insights, we would like to develop future probiotics studies.

## Acknowledgements

We thank Satoru Furukawa of the FRC for his technical advice, and Kayo Yasuda of Tokai University, Youko Yasutomi of Tokushima University, Kyoko. Takada, and Kaoru Yoshitaka of Tokyo University of Agriculture for their technical assistance. This work was supported by grants from the Japanese Ministry of Education, Culture, Sports, Science and Technology.

## Supporting information

**S1 Fig. Evaluation of the lifespan of *C. elegans Δfer-15.***

*E. coli* OP50 (gray), *S. thermophiles* NRIC0256T(green), *L. plantarum* P1-2 (red), and *P. pentosaceus* Be1 (blue) were administered to *C. elegans Δfer-15* when the growth stage reached L4. The mutants were hatched on pH stat GYP medium, and their lifespan was monitored until annihilation. Statistical analysis was carried out by Student’s t-test and Tukey’s multiple-range test. The least significant difference test was used for means separation at *P* < 0.05 within each strain. One hundred animals were measured for each strain at 25°C.

**S2 Fig. Administration effect of *P. pentosaceus* Be1 strain exhibiting hydrogen peroxide eliminating activity on iron-overloaded rats: a colonic mucosal lipid peroxidation model.**

*P. pentosaceus* Be1 was administered to iron-overloaded rats, and the MDA levels in the colonic mucosa were compared to those of the healthy (control) and iron-overloaded rats (Fe). *L. plantarum* P1-2 and *S. thermophilus* NRIC0256^T^ were also tested as the control strain. The data are the mean values ± SD (n = 4).

